# Distinct 5′ and 3′ Coverage Biases Shape Transcriptome Interpretation in Nanopore Direct RNA versus PCR-cDNA Sequencing

**DOI:** 10.1101/2025.10.13.681987

**Authors:** Rebecca E Lane, Eleanor Calcutt, Anandagopal Srinivasan, Vicki Gamble, Udo Oppermann, Jianfeng Sun, Adam P Cribbs

## Abstract

Long-read RNA sequencing enables isoform-resolved transcriptomics, but library preparation introduces systematic biases that shape biological interpretation. We benchmarked Oxford Nanopore’s two protocols—PCR-cDNA and direct RNA—using SKMM2 myeloma cells stimulated with interleukin-6 (IL-6) and ERCC synthetic spike-ins. Direct RNA produced longer, higher-quality reads and more high-confidence isoforms, but showed pronounced 5′ coverage loss. PCR-cDNA yielded shorter fragments with 3′ underrepresentation, detecting more low-abundance transcripts at reduced confidence. Protocol-specific biases had major consequences: differential expression analysis revealed limited overlap in IL-6–responsive genes, and pathway enrichment was broader in direct RNA. At the isoform level, differential transcript usage was almost entirely protocol-specific, with case studies (e.g. RPL22L1, GRB2, RNF220) illustrating concordance and divergence. ERCC controls confirmed these biases as technical rather than biological. Together, our results show that while both methods provide accurate gene-level quantification, transcript-level conclusions depend critically on protocol choice, highlighting the need for careful selection in long-read transcriptomics.

## Introduction

Long-read sequencing technologies have transformed transcriptomic research by enabling direct measurement of full-length RNA molecules and their isoforms(1). Among these, Oxford Nanopore Technologies (ONT) has developed protocols capable of sequencing either cDNA molecules generated from reverse-transcribed RNA(2, 3), or native RNA molecules themselves(4-8). These two approaches—PCR-cDNA and direct RNA sequencing—differ markedly in library preparation, sequencing chemistry, throughput, and bias profiles. Understanding these differences is critical for researchers seeking accurate transcriptome characterisation.

The ONT PCR-cDNA protocol benefits from relatively low input requirements, high throughput, and robust detection of low-abundance transcripts. However, the multiple enzymatic steps required—reverse transcription, strand-switching, ligation and PCR amplification—introduce artefacts(9). These include preferential amplification of shorter molecules, underrepresentation of GC-rich regions, and incomplete reverse transcription leading to truncated reads. Such biases are well documented in short-read cDNA sequencing and remain a challenge for nanopore-based approaches (10, 11).

Direct RNA sequencing eliminates the need for PCR amplification, allowing direct sequencing of native RNA molecules. This provides several advantages: removal of PCR-induced artefacts, improved isoform resolution through long contiguous reads, and the unique ability to detect base modifications such as N6-methyladenosine(12, 13). However, these benefits come with trade-offs, including higher input requirements, lower throughput, multiplexing restrictions and reduced read accuracy compared to PCR-cDNA.

Benchmarking studies have demonstrated that both protocols provide accurate gene-level quantification (14), but their biases diverge at the transcript and isoform levels (15). Consequently, biological inferences—particularly regarding alternative splicing, isoform switching, or differential transcript usage—may depend heavily on the chosen protocol.

In this study, we perform a systematic comparison of Oxford Nanopore Technologies (ONT) PCR-cDNA and direct RNA sequencing protocols. We first benchmark both methods in the SKMM2 multiple myeloma cell line stimulated with interleukin-6 (IL-6), providing a controlled model to evaluate how protocol-specific biases impact gene expression, isoform detection, and pathway-level interpretation. This analysis reveals areas of concordance—such as gene-level quantification—and key points of divergence, particularly in transcript isoform usage and differential expression outcomes. To validate that these biases arise from technical rather than biological sources, we extended our evaluation using ERCC synthetic spike-in controls. These confirmed differences in sensitivity, coverage uniformity, and quantitative accuracy between the two protocols. Together, our findings establish a technical framework for understanding the strengths and limitations of ONT RNA sequencing strategies and for guiding protocol choice long-read in transcriptomic studies.

## Results

### Protocol-dependent read characteristics and transcript detection in SKMM2 myeloma cells

To assess protocol-specific differences in nanopore transcriptomics, we first compared PCR-cDNA and direct RNA sequencing in SKMM2 cells with or without IL-6 stimulation (Figure 1). Direct RNA sequencing consistently generated longer reads, with median read lengths of ∼800 bp compared to ∼500 bp for PCR-cDNA (Fig. 1a). Read length distributions revealed distinct profiles: PCR-cDNA was enriched for shorter fragments (∼500 bp peak), whereas direct RNA yielded a broader distribution with greater representation of longer molecules (Fig. 1b). Read quality scores were slightly higher for direct RNA (median 14–15) compared to PCR-cDNA (median 13– 14), with no marked influence of IL-6 treatment (Fig 1c).

**Figure 1.**
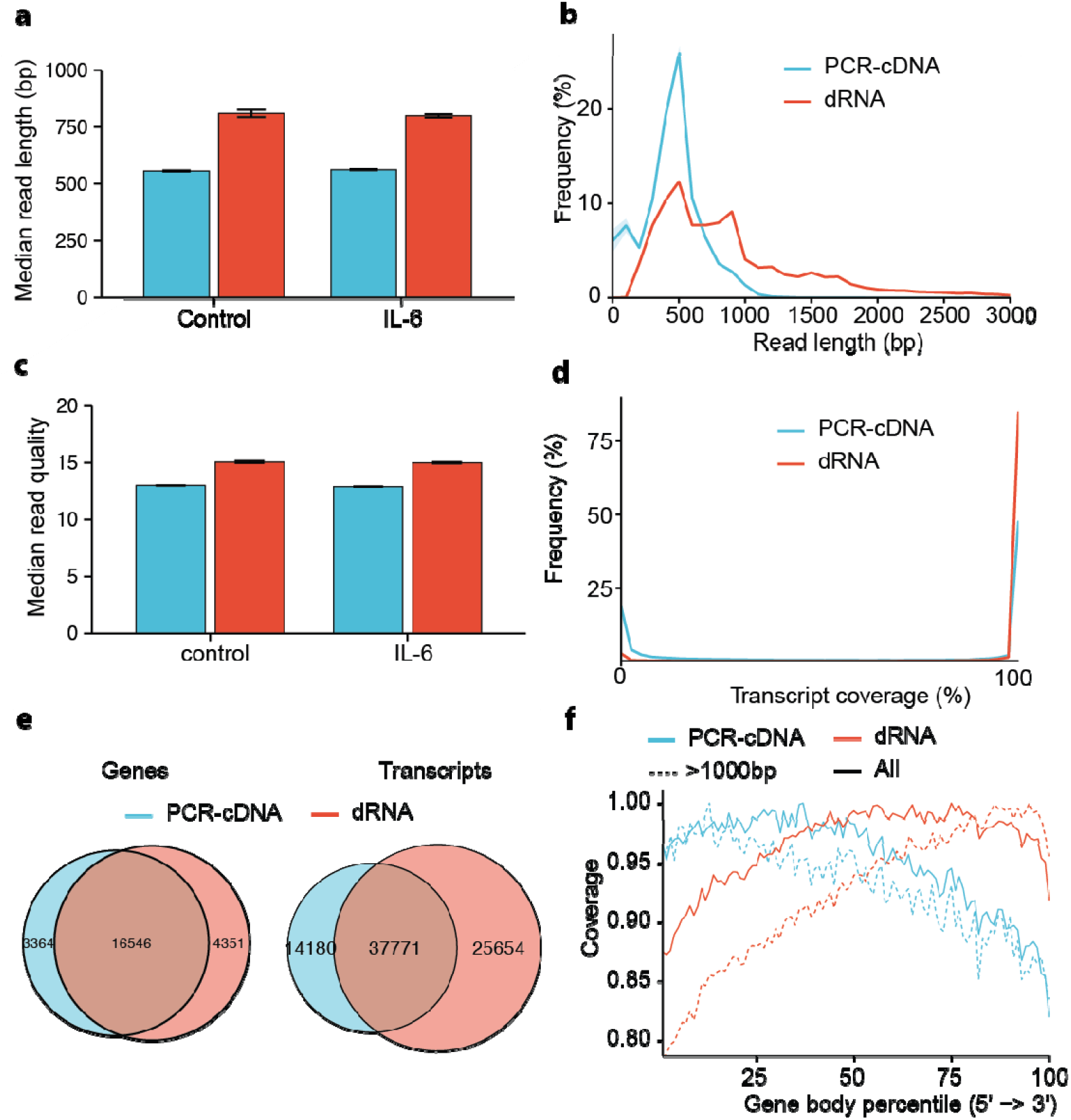
Read characteristics and transcriptome coverage differ between PCR-cDNA and direct RNA sequencing. (a) Median read length across biological replicates of SKMM2 myeloma cells cultured with or without IL-6 stimulation. Direct RNA (dRNA, red) yielded consistently longer reads than PCR-cDNA (blue). (b) Distribution of read lengths, showing enrichment of short fragments in PCR-cDNA and broader length representation in direct RNA. (c) Median read quality scores by protocol, with direct RNA achieving slightly higher values. (d) Transcript coverage profiles, highlighting 5′ coverage loss in direct RNA and 3′ underrepresentation in PCR-cDNA. (e) Venn diagrams of gene- and transcript-level detection, showing ∼16,500 genes shared, with PCR-cDNA enriched for unique low-abundance genes and direct RNA identifying more isoforms. (f) Normalised coverage along transcript bodies, demonstrating protocol-specific biases: PCR-cDNA underestimates coverage towards the 3′ end, while direct RNA shows loss of coverage at the 5′ end.

Coverage profiling across transcripts highlighted characteristic biases of each protocol. PCR-cDNA showed progressive loss of coverage towards the 3′ end of coding sequence, consistent with incomplete reverse transcription and amplification inefficiency. In contrast, direct RNA exhibited reduced coverage at the 5′ end, reflecting the 3′–5′ sequencing orientation and RNA degradation (Fig. 1d and Fig. 1e).

These technical biases translated into differences in transcript detection, despite normalising for read depth across both protocols. At the gene level, both methods recovered a comparable number of genes, with ∼16,500 shared and smaller sets uniquely identified by PCR-cDNA (8364) or direct RNA (4351) (Fig 1f). At the transcript level, however, differences were more pronounced: direct RNA identified over 63,000 isoforms, whereas PCR-cDNA identified ∼51,000, with ∼37,700 shared between protocols. This suggests that direct RNA sequencing may be better able to support isoform-level inference, although differences between the sequencing methods suggest biases within each sequencing approach.

### Transcript detection is strongly protocol-dependent

We next compared the breadth and confidence of transcript detection between protocols. Direct RNA sequencing (dRNA) identified a greater number of transcripts than PCR-cDNA, with ∼63,425 versus ∼51,951 detected, and ∼37,771 shared between methods (Fig. 2a–b). This broad recovery by dRNA was particularly evident at higher confidence levels. At high confidence, dRNA uniquely detected 12,645 transcripts compared to only 753 for PCR-cDNA. At medium confidence, both protocols captured ∼10,743 shared isoforms, but dRNA contributed more unique transcripts (25,174 vs 14,663). In contrast, at low confidence PCR-cDNA recovered a much larger fraction of unique transcripts (16,728 vs 5,799), reflecting increased sensitivity to low-abundance fragments but at reduced assignment certainty.

**Figure 2.**
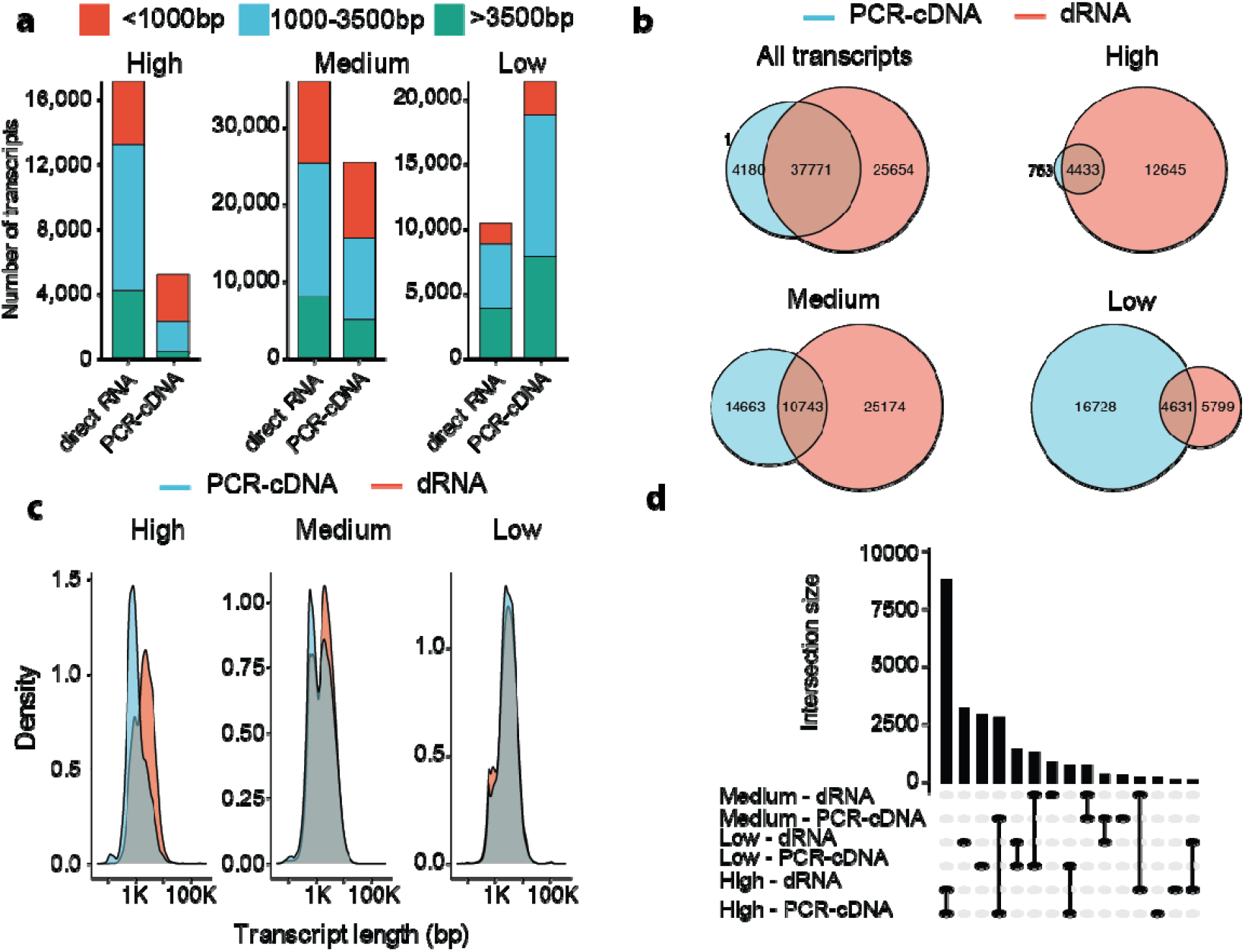
Transcript detection differs by protocol, confidence, and length. (a) Number of transcripts detected by PCR-cDNA (blue) and direct RNA (dRNA, red) across high-, medium-, and low-confidence categories, stratified by transcript length (<1000 bp, 1000–3500 bp, >3500 bp). PCR-cDNA is enriched for shorter, low-confidence transcripts, whereas direct RNA recovers a greater proportion of longer isoforms. (b) Venn diagrams of transcript detection across protocols. While 37,771 transcripts were shared, direct RNA uniquely identified >25,000 isoforms, particularly at high and medium confidence. PCR-cDNA contributed more unique transcripts at low confidence. (c) Transcript length distributions for high-, medium-, and low-confidence sets. PCR-cDNA libraries are biased towards short isoforms, while direct RNA captures a broader size distribution. (d) Intersection analysis of transcripts across confidence categories and protocols. Direct RNA shows enhanced recovery of high- and medium-confidence isoforms, while PCR-cDNA contributes disproportionately to low-confidence calls.

Transcript length distributions further highlighted protocol-specific biases (Fig. 2c). PCR-cDNA libraries were strongly skewed toward short isoforms (<1000 bp), particularly at low confidence, consistent with amplification-driven overrepresentation of short molecules. Direct RNA sequencing, in contrast, recovered a more balanced spectrum of isoform sizes, including robust detection of long transcripts (>3500 bp).

Intersection analysis confirmed these trends (Fig. 2d). dRNA preferentially contributed high- and medium-confidence isoforms, while PCR-cDNA disproportionately inflated the low-confidence category. Together, these results demonstrate complementary detection properties: PCR-cDNA maximises sensitivity at the expense of confidence and length balance, whereas direct RNA sequencing provides broader and more reliable identification of high-confidence, long isoforms.

### Divergent IL-6–induced transcriptional responses across protocols

To assess how protocol choice influences biological interpretation, we compared IL-6–stimulated SKMM2 multiple myeloma cells using both PCR-cDNA and direct RNA sequencing. Although both approaches detected IL-6–responsive changes, the breadth and concordance of results differed markedly (Fig. 3).

**Figure 3.**
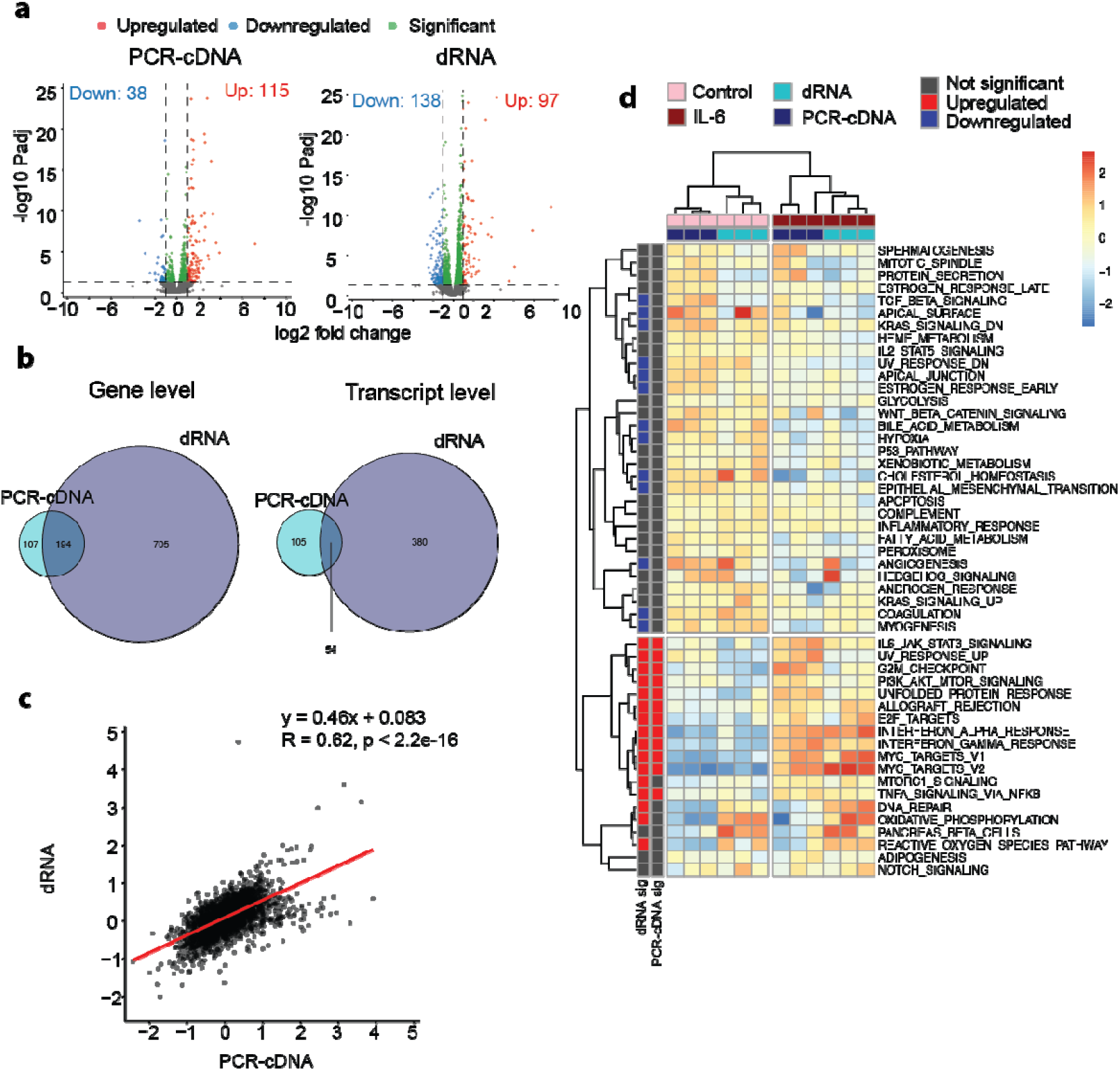
Divergent IL-6–induced transcriptional responses detected by PCR-cDNA and direct RNA sequencing. (a) Differential expression analysis of SKMM2 cells stimulated with IL-6. Volcano plots show significantly upregulated (red) and downregulated (blue) genes for PCR-cDNA (left, 115 up, 38 down) and direct RNA (right, 97 up, 138 down). (b) Venn diagrams at the gene (left) and transcript (right) levels. Overlap between protocols is limited, with only 194 genes and 54 transcripts identified by both, while the majority of differentially expressed features are protocol-specific. (c) Correlation of fold-change estimates between protocols. While overall correlation was moderate (R = 0.62), PCR-cDNA compressed fold-change dynamic range relative to direct RNA. (d) Pathway-level analysis of differentially expressed genes. Heatmap of hallmark gene set enrichment reveals broader pathway activation with direct RNA, including IL-6/JAK–STAT signalling, MYC targets, and inflammatory programmes, whereas PCR-cDNA highlights fewer significantly perturbed pathways.

At the differential expression level, PCR-cDNA sequencing identified 301genes, whereas direct RNA sequencing detected substantially more, with 899 genes (Fig. 3a). Overlap between the two approaches was modest: only 194 differentially expressed genes were shared, with the majority uniquely detected by one protocol. This divergence was even more pronounced at the transcript level, where only 54 isoforms were shared, and most differentially expressed transcripts were protocol-specific (Fig. 3b).

Direct comparison of log□ fold-change values showed only moderate correlation between protocols (R = 0.62; Fig. 3c), with PCR-cDNA compressing the dynamic range relative to direct RNA. This compression likely contributes to under-detection of isoform-specific responses.

Downstream pathway analysis further highlighted protocol-specific biases (Fig. 3d). PCR-cDNA sequencing reported a restricted set of enriched pathways, whereas direct RNA sequencing revealed broader IL-6–responsive signatures, including robust activation of JAK/STAT3 signalling, MYC target genes, and inflammatory programmes, alongside repression of metabolic and epithelial pathways.

Together, these results demonstrate that while both ONT protocols detect IL-6– induced transcriptional responses, the scope, sensitivity, and inferred biology differ substantially, with direct RNA sequencing providing broader and more nuanced coverage of isoform- and pathway-level responses.

### Protocol-specific detection of differential transcript usage

To further dissect protocol-specific effects, we compared differential transcript usage (DTU) across selected genes (Fig. 4). These examples illustrate both concordance and divergence in isoform-level interpretation between PCR-cDNA and direct RNA sequencing.

**Figure 4.**
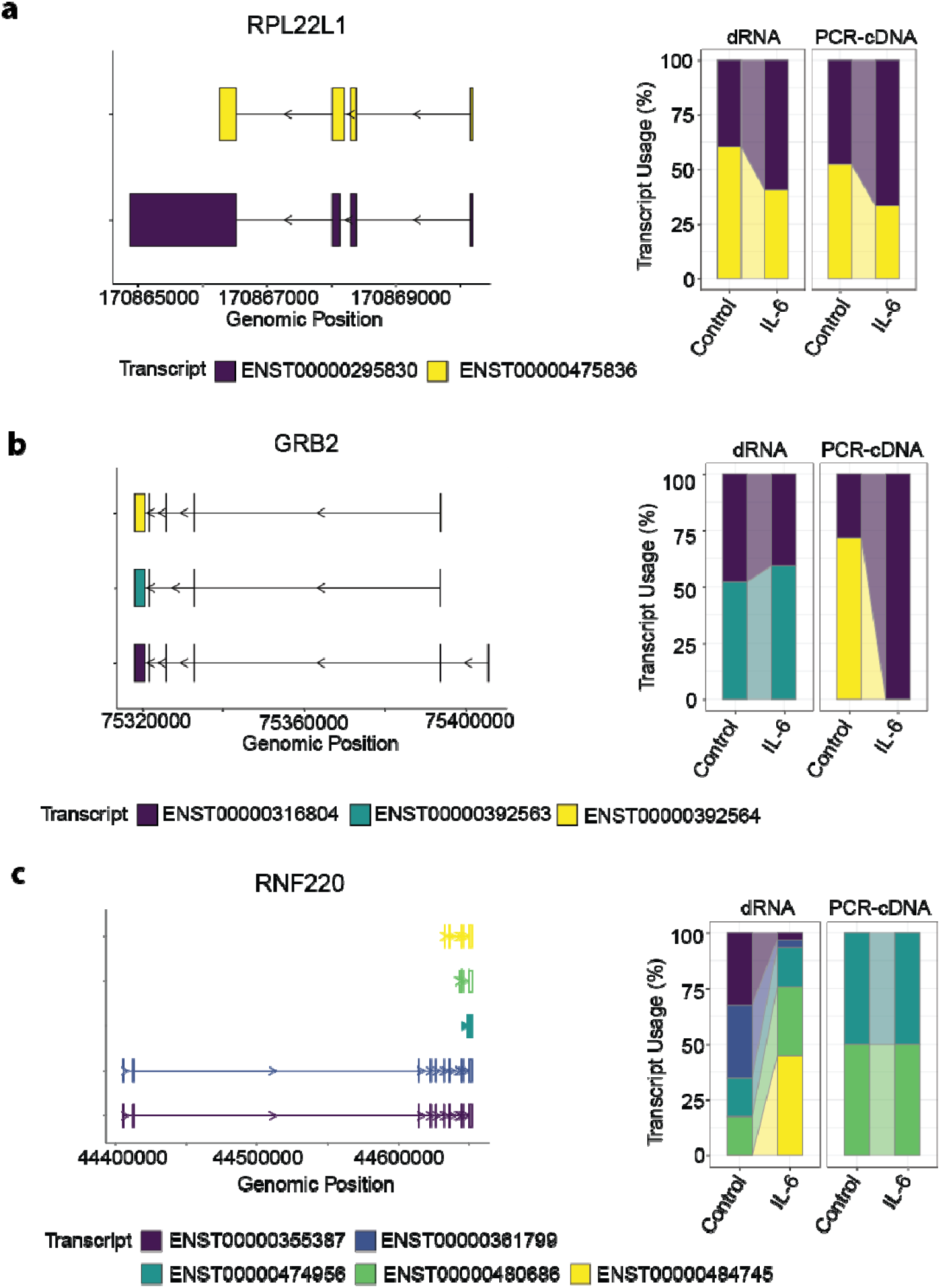
Isoform-specific responses reveal protocol-dependent concordance and divergence. (a) RPL22L1: Two isoforms were detected across conditions. Both PCR-cDNA and direct RNA identified shifts in isoform usage upon IL-6 stimulation, with broadly concordant patterns. (b) GRB2: Three isoforms were detected. While both protocols captured IL-6– induced changes, the relative contribution of minor isoforms differed between PCR-cDNA and direct RNA. (c) RNF220: Five isoforms were identified. Direct RNA resolved multiple isoforms with distinct usage patterns, whereas PCR-cDNA predominantly supported two transcripts, underestimating isoform diversity.

For RPL22L1, both protocols consistently identified a shift in isoform usage upon IL-6 stimulation, supporting concordant detection of DTU (Fig 4a). This indicates that for certain highly expressed genes with relatively simple isoform structures, both methods provide reproducible isoform-level inferences.

By contrast, GRB2 showed a striking divergence: PCR-cDNA suggested a strong switch from ENST00000392563.5 to ENST00000316804.10, whereas direct RNA sequencing supported the continued dominance of ENST00000316804.10 with limited evidence for switching (Fig. 4b). This discrepancy may arise from PCR amplification bias against longer isoforms.

A different pattern was observed for RNF220, where direct RNA sequencing uniquely identified IL-6–induced changes in isoform proportions, whereas PCR-cDNA showed little evidence of DTU (Fig. 4c). This reflects direct RNA’s greater capacity to capture long and complex isoforms with higher confidence.

Taken together, these case studies demonstrate that while gene- and pathway-level signals are broadly robust, transcript-level inferences are strongly shaped by the choice of sequencing protocol. The minimal overlap in DTU calls underscores the need for caution when drawing isoform-specific biological conclusions from single-protocol datasets.

### Novel transcript discovery reveals further protocol-specific divergence

We next examined the ability of PCR-cDNA and direct RNA sequencing to detect and quantify novel isoforms (Fig. 5). Principal component analysis (PCA) showed clear separation by both protocol and treatment, indicating that technical biases and IL-6 stimulation both contribute to the observed variance (Fig. 5a). Differential expression analysis of novel transcripts identified by bambu revealed 665 candidates, with several highly significant isoforms upregulated in IL-6–treated cells (Fig. 5b). Heatmap clustering highlighted distinct expression patterns, with direct RNA capturing a broader repertoire of novel transcripts compared to PCR-cDNA (Fig. 5c). Intersection analysis confirmed this trend: direct RNA identified the largest set of unique novel isoforms, while overlap between protocols was limited (Fig. 5d). These results demonstrate that protocol choice not only influences detection of annotated isoforms but also strongly affects the discovery and quantification of novel transcripts.

**Figure 5.**
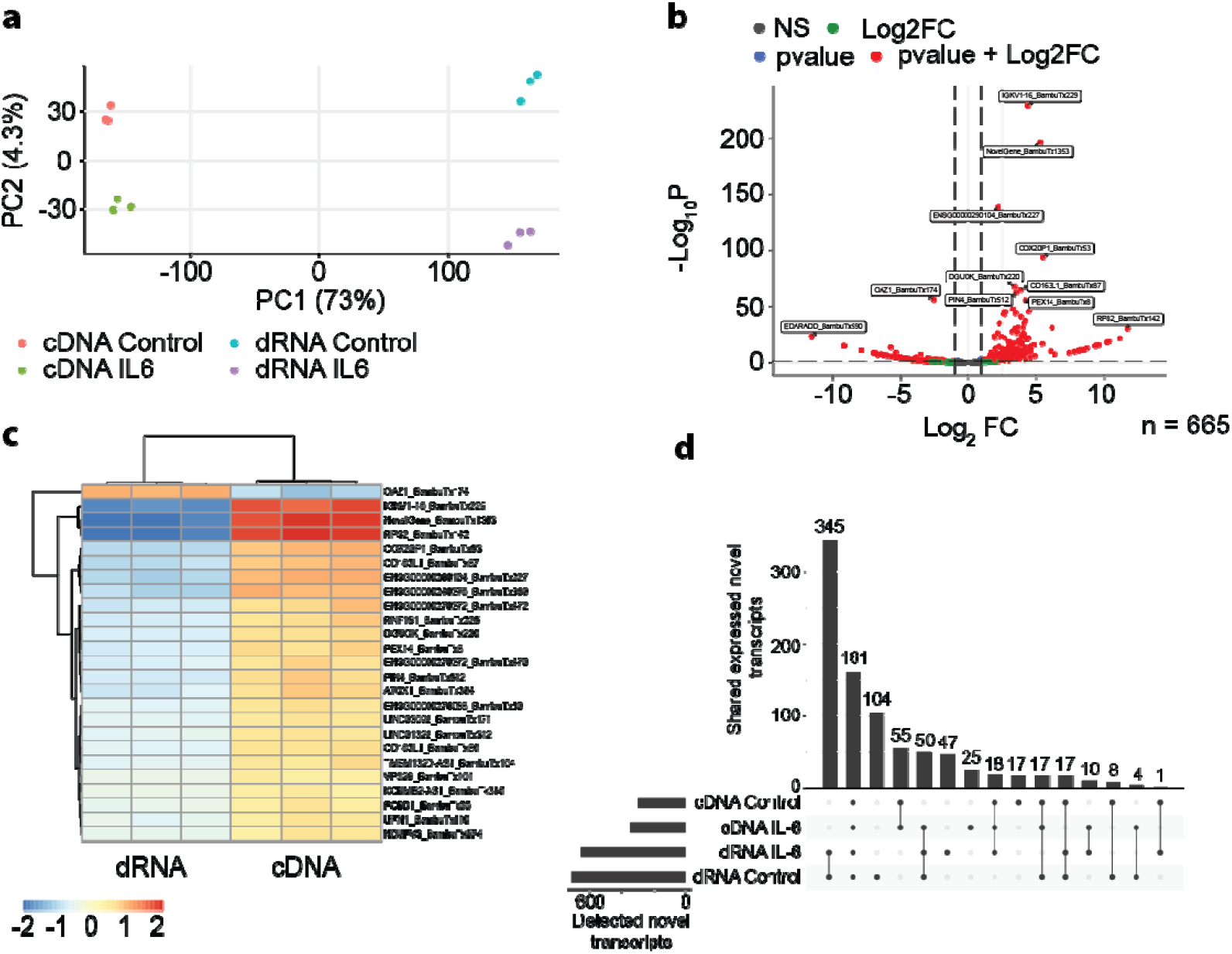
Principal component analysis, differential expression, and detection of novel transcripts across protocols. (a) Principal component analysis (PCA) of SKMM2 cells sequenced by PCR-cDNA or direct RNA (dRNA) protocols, with or without IL-6 stimulation. Clear separation by protocol and treatment indicates protocol-specific bias as well as biological response. (b) Differential expression analysis of novel transcripts detected by bambu. Volcano plot shows significant upregulated (red) and downregulated (green) transcripts after IL-6 treatment, with labelled examples of highly significant candidates. (c) Heatmap of the top differentially expressed novel transcripts, illustrating distinct expression patterns across protocols and treatment conditions. (d) Intersection analysis of novel transcript detection across conditions. Bar plots indicate the number of novel transcripts detected per protocol and treatment, while the Upset plot shows shared and unique subsets. Direct RNA sequencing identified a larger set of novel isoforms compared with PCR-cDNA, with limited overlap between protocols.

### Synthetic ERCC benchmarking validates protocol-specific biases

To disentangle biological variability from protocol artefacts, we sequenced ERCC synthetic spike-in controls using both PCR-cDNA and direct RNA protocols (Fig. 6). At the gene level, both protocols produced highly accurate quantification, with transcript abundances strongly correlated with expected concentrations (R = 0.97 for PCR-cDNA; R = 0.98 for direct RNA). Root mean square error (RMSE) and mean absolute error (MAE) values were lower for direct RNA (RMSE = 0.68, MAE = 0.47) compared with PCR-cDNA (RMSE = 0.86, MAE = 0.66), confirming the higher quantitative accuracy of direct RNA (Fig. 6a).

**Figure 6.**
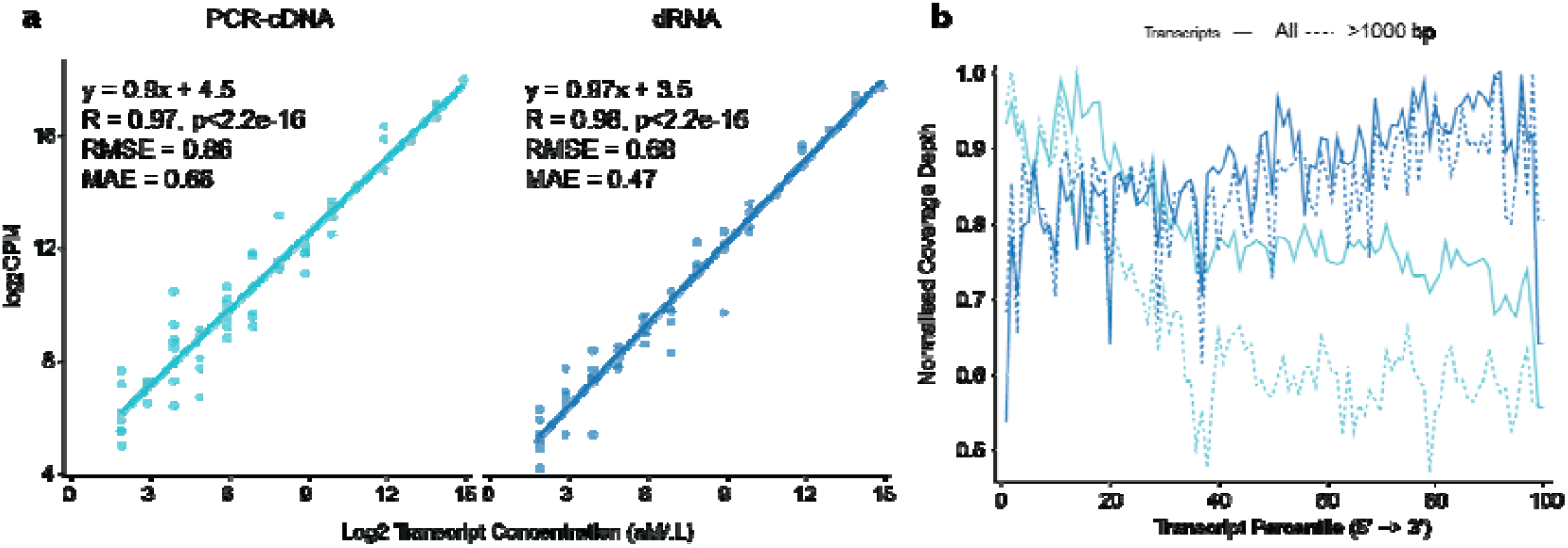
ERCC spike-in controls validate protocol-specific biases. (a) Correlation of measured expression (log□ CPM) with known ERCC transcript concentrations for PCR-cDNA (left) and direct RNA (dRNA, right). Both protocols achieved strong correlations (R = 0.97 for PCR-cDNA, R = 0.98 for direct RNA), but direct RNA showed lower root mean square error (RMSE = 0.68) and mean absolute error (MAE = 0.47) compared with PCR-cDNA (RMSE = 0.86, MAE = 0.66), indicating higher quantitative accuracy. (b) Normalised coverage profiles of ERCC transcripts reveal protocol-specific biases consistent with cellular transcriptomes. PCR-cDNA libraries show underrepresentation towards the 3′ end, whereas direct RNA exhibits reduced coverage at the 5′ end. These effects are observed across all spike-ins and are more pronounced in longer transcripts (>1000 bp).

Coverage profiles, however, recapitulated the characteristic biases of each protocol (Fig. 6b). PCR-cDNA showed progressive coverage loss at the 3′ ends of spike-ins, consistent with incomplete reverse transcription and PCR artefacts. Conversely, direct RNA sequencing displayed reduced coverage at the 5′ ends, reflecting sequencing orientation and RNA degradation. These biases were consistent with those observed in SKMM2 myeloma cells, validating them as intrinsic to the protocols rather than driven by biological features.

Together, the ERCC benchmarking confirms that while both methods provide accurate abundance estimates at the gene level, protocol-specific biases in coverage and isoform detection systematically shape transcriptome interpretation.

## Discussion

This study highlights both the complementary strengths and the limitations of ONT PCR-cDNA and direct RNA sequencing when applied to isoform-resolved transcriptomics. Using SKMM2 multiple myeloma cells as a controlled model, we demonstrate that while both protocols capture IL-6–induced transcriptional responses, they diverge substantially at the isoform and transcript level. Direct RNA sequencing consistently yielded longer reads, more balanced transcript coverage, and higher-confidence isoform assignments, whereas PCR-cDNA provided greater sensitivity for low-abundance transcripts but at the cost of amplification bias and reduced length representation. These protocol-specific biases were independently validated using ERCC synthetic spike-ins, which confirmed that PCR-cDNA libraries tend to enrich for shorter fragments and favour 5′ coverage, while direct RNA libraries underrepresent 5′ regions, consistent with RNA degradation.

A further source of technical bias arises during the reverse transcription step of PCR-cDNA library preparation. Incomplete reverse transcription can lead to 3′ truncation of cDNA molecules, which in turn may partially or completely remove the unique molecular identifier (UMI) sequence. As we and others have shown previously (9, 16, 17), this truncation introduces a significant bioinformatic challenge, impeding the accurate identification and collapsing of reads derived from the same RNA molecule. Reads with truncated or unrecognisable UMIs are frequently discarded, resulting in the under-representation of valid long molecules in the final dataset. This not only reduces the accuracy of transcript quantification but also diminishes statistical power for downstream analyses. Such effects are a plausible mechanistic explanation for the weaker identification of differentially expressed genes by the PCR-cDNA method compared to direct RNA sequencing.

A notable consequence of these biases is their impact on biological interpretation. At the gene and pathway levels, both protocols recovered canonical IL-6–responsive signatures such as JAK/STAT3 and MYC targets, though direct RNA consistently detected broader sets of differentially expressed genes and pathways. At the isoform level, however, overlap was minimal: differential transcript usage (DTU) calls were largely protocol-specific, and novel transcript discovery diverged between approaches. These findings emphasise that transcript-level biology inferred from long-read sequencing may reflect protocol-specific technical artefacts as much as genuine biology.

Our findings align with and extend two recent benchmarking studies. Pardo-Palacious et al. (15) compared direct RNA and cDNA approaches and highlighted challenges in transcript isoform detection, quantification and de novo transcript detection, with direct RNA sequencing but also emphasised its lower sensitivity. Similarly, Chen et al. underscored protocol-specific biases in coverage, with direct RNA limited by throughput and input requirements(14). Both studies, like ours, stress that while gene-level quantification is broadly concordant between protocols, isoform-level interpretation is highly protocol-dependent. Importantly, our work adds a direct biological case study—IL-6 signalling in myeloma—showing how these technical biases directly propagate into differential expression, pathway enrichment, and isoform usage analyses.

Pragmatically, protocol choice should be guided by study aims. For isoform discovery and differential transcript usage, direct RNA sequencing offers higher confidence despite greater RNA input and low throughput. For experiments prioritising sensitivity to low-abundance transcripts or when input is limiting, PCR-cDNA remains valuable. Taken together with prior benchmarking efforts, our analysis provides a technical framework for selecting ONT sequencing protocols and for interpreting their results with appropriate caution, particularly when drawing biological conclusions from isoform-level data.

## Methods

### SKMM2 Cell Culture and RNA Extraction

SKMM2 cells (DSMZ: ACC 430) were cultured in Gibco™ RPMI 1640 medium (11530586, Fisher Scientific, UK), supplemented with 10% FBS (SV30160.03, Cytiva Life Sciences, UK), and maintained at 37°C in a 5% CO□ atmosphere. Cells were seeded into 12-well plates in triplicate and treated with either 1 ng/mL interleukin-6 (IL-6) in 0.2% BSA or with 0.2% BSA alone for 18 hours. Following treatment, cells were harvested and cell pellets resuspended in 350 µL Trizol reagent. RNA extraction was performed using the Zymo Direct-zol RNA Miniprep kit (R2051, Zymo Research, USA) according to the manufacturer’s protocol, with RNA eluted in 25 µL DNase/RNase-free water. RNA concentration was determined using a NanoPhotometer NP80 UV/Vis Spectrophotometer (Implen, Germany), while RNA integrity and size profile were assessed using the Agilent 4150 TapeStation System (Agilent Technologies, UK) with an RNA ScreenTape (Agilent Technologies, UK). All RNA samples had a RNA Integrity Number (RIN) between 9.7 and 9.8.

### PCR-cDNA Library Preparation and Sequencing

Library preparation was performed using the PCR-cDNA Barcoding Kit (SQK-PCB111.24, Oxford Nanopore Technologies, UK) following the manufacturer’s instructions. A total of 200 ng of RNA (IL-6 treated, n=3; untreated, n=3) or 16 ng of ERCC RNA Spike-In Mix 1 (Thermo Fisher, UK) was used for cDNA synthesis. For SKMM2 samples, 14 cycles of PCR amplification were conducted, while 16 cycles were used for the ERCC mix. Library concentrations were quantified using a Qubit 2.0 Fluorometer (Thermo Fisher, UK) with the Qubit dsDNA HS Assay kit (Q32851, Thermo Fisher, UK). Library quality was assessed on a 4150 TapeStation System (Agilent Technologies, UK) using a high sensitivity D5000 ScreenTape. For sequencing, 2 fmol of each barcoded library were pooled, and 12 libraries (24 fmol) were loaded onto a PromethION R9.4.1 flow cell (FLO-PRO002, Oxford Nanopore Technologies, UK). Sequencing was performed on a PromethION 24 instrument (Oxford Nanopore Technologies, UK) using MinKNOW software (v24.06.15), with live base-calling using the Dorado high accuracy model (v7.4.14). The sequencing run lasted for 72 hours, with a minimum Q score of 9. For the ERCC mix, 5 fmol of the prepared library was loaded onto an R10.4.2 Flongle flow cell (FLO-FLG114, Oxford Nanopore Technologies, UK), and sequencing was conducted on a MinION Mk1B instrument with the Flongle adapter (Oxford Nanopore Technologies, UK). Live base-calling was performed using the Dorado high accuracy model (v7.3.11) with a 24-hour run time, a minimum read length of 20 bp, and a minimum Q score of 9.

### Direct RNA Library Preparation and Sequencing

Library preparation was performed using the Direct RNA Sequencing Kit (SQK-RNA004, Oxford Nanopore Technologies, UK) following the manufacturer’s instructions. A total of 1 µg of RNA (IL-6 treated, n=3; untreated, n=3) or ∼150 ng of ERCC RNA Spike-In Mix 1 (n=1) was used for direct RNA sequencing. The concentration of the prepared libraries was measured using a Qubit 2.0 Fluorometer (Thermo Fisher, UK) and the Qubit dsDNA HS Assay kit (Q32851, Thermo Fisher, UK). All libraries exceeded the recommended 30 ng yield. Library size profiles were assessed using the Agilent 4150 TapeStation System (Agilent Technologies, UK) with a high sensitivity D5000 ScreenTape. Libraries were loaded in their entirety onto separate PromethION RNA Flow Cells (FLO-PRO004RA, Oxford Nanopore Technologies, UK). Sequencing was performed on a PromethION 24 instrument (Oxford Nanopore Technologies, UK) with MinKNOW software (v24.06.15). Live base-calling was performed using the Dorado RNA high accuracy model (v7.4.14). Sequencing lasted for 72 hours, with a minimum read length of 200 bp and a minimum Q score of 9.

### Data processing and analysis

Reads passing quality filters were demultiplexed by MinKNOW software, where applicable, and concatenated into a single FASTQ file for each sample. For PCR-cDNA data, adapter trimming and read orientation were performed using pychopper (v2.7.10, Oxford Nanopore Technologies, UK). Reads shorter than 50 bp were discarded. Trimmed reads were aligned to the hg38 human genome assembly (GRCh38.p14, GENCODE v46) using minimap2 (v2.28) with spliced alignment mode, a k-mer size of 14, a minimum primary-to-secondary score ratio of 0.9, and canonical splice-site searching on the forward strand only. Direct RNA sequencing reads were aligned similarly, without adapter trimming. For ERCC reference transcripts, transcript quantification was performed using featureCounts within the Subread package (v2.0.6). Gene and transcript count matrices for SKMM2 cell line samples were generated using bambu (v3.6.0). Transcript coverage depth was calculated using the gene body coverage module of RSeQC (v5.0.4). All subsequent analyses and data visualisation were performed in R (version 4.4.1).

Differential gene expression was analysed using DESeq2 (v1.44.0) with a false discovery rate (FDR) of 5%. Count normalization was performed using variance stabilizing transformation (VST), and VST-normalized data were used for visualisation and exploratory analyses. Gene Set Variation Analysis (GSVA) was performed using the GSVA package (v2.0.0) with hallmark gene sets from the Molecular Signatures Database (MSigDB). Differential expression of gene sets was assessed using linear modelling with empirical Bayes moderation (FDR = 5%).

Prior to testing for differential transcript usage (DTU), genes and transcripts were filtered using DRIMSeq (v1.52.0). Genes were required to be expressed in all samples at >5 counts, and transcripts had to be expressed in >3 samples, with >5 counts and >10% transcript usage for the gene. DTU analysis was conducted using DEXSeq (v1.52.0), with FDR set to 5% for both gene-wise and transcript-wise analyses. Stage-wise adjustment for DTU was performed using StageR (v1.28.0) with FDR set to 5%, as described by van den Berge et al. Graphics were produced using ggplot2 (v3.5.1), ggpubr (v0.6.0), ggvenn (v0.1.10), ggtranscript (v1.0.0), ggalluvial (v0.12.5), and pheatmap (v1.0.12).

## Data availability

Sequencing data have been deposited to the Gene Expression Omnibus under the accession number GSE306225. All analysis was performed using hg38 ensembl 101 version.

## Code availability

All custom code used within this analysis are available on GitHub (https://github.com/cribbslab/ONT_comparison)

## Author contributions

R.E.L performed analysis, data curation and drafting of the manuscript. E.C performed sequencing, quality control and manuscript editing. A.S. performed analysis and manuscript editing. V.G supported experimental work including cell culture and RNA preparation. U.O. and J.S. contributed study supervision and manuscript review. A.P.C conceived and supervised the project, acquired funding, administered the study, and led manuscript revision as corresponding author. All authors approved the final manuscript

